# Gain and loss of gut symbionts during stingless bee diversification is linked to host group size

**DOI:** 10.64898/2026.01.16.699994

**Authors:** Nickole A. Villabona, Alejandro Vazquez, Jennifer Schlauch, Claus Rasmussen, Richard Joyce, Christopher M. Martinez, Tobin J. Hammer

## Abstract

Microbiomes typically vary between closely related host species, but how and why they diverge during host diversification remain poorly understood. In bees, the evolution of eusociality is thought to have promoted symbiosis with specialized gut bacteria, including *Snodgrassella* and *Gilliamella*. While these symbionts are common in bumble bees and honey bees, some stingless bees, a closely related clade of eusocial bees, lack them. Here, we tested whether group size, a fundamental trait of social organisms with implications for microbial transmission and function, explains symbiont gain and loss in stingless bees. Using specimens from a global (though mostly Neotropical) collection, we performed 16S rRNA gene sequencing and qPCR on adult worker bees representing 29 species from 18 genera. Phylogenetic models indicate that bee species with larger colonies are more likely to harbor *Snodgrassella* and *Gilliamella*. We propose two underlying mechanisms: challenges faced by large groups (e.g., disease risk) select for symbionts that supply needed functionality (e.g., pathogen protection), or large groups are less prone to transmission bottlenecks and stochastic symbiont loss. While deeper sampling within species is still needed, these findings highlight group size as a driver of microbiome evolution in stingless bees and, potentially, in other social hosts.

## Introduction

One of the most commonly observed ecological patterns in host-microbiome research is variation in microbial community composition among host species [1–5]. From an evolutionary perspective, this variation may reflect a dynamic history of gains and losses of particular microbiome members (symbionts) during host diversification [6]. Understanding what drives these dynamics is important, as microbial symbionts often play key roles in host physiology, immunity, and other functions [7–9]. Among animals, divergence or convergence in diet often explains a substantial fraction of gut microbiome variation across species [1,2,10–13]. However, animals vary not only in what they eat, but also in their behavior and physiology, and the influence of these non dietary traits on gut microbiome evolution remains poorly understood.

Social behavior has evolved numerous times in animals and has shaped multiple aspects of their biology [14], including the gut microbiome [15–17]. Most studies on the “social microbiome” [18] focus on associations between social traits and the microbiome within species over ecological time scales [15,19]. Individuals from the same social group often share more similar gut microbiomes than those from different groups, even after controlling for diet, genetics, and habitat, suggesting that social interactions can facilitate microbial exchange and persistence [15,16,19]. Conversely, laboratory experiments have shown that the gut microbiome can affect social traits [20,21]. However, less is known about how social behavior and the gut microbiome may influence one another over macroevolutionary time scales.

Bees present an example of the role of sociality in shaping gut microbiome evolution. Modes of sociality range from complex eusociality in the corbiculate Apinae (honey bees, bumble bees, and stingless bees), to facultative or simple social groups, and to solitary life histories [22–24]. Despite the fact that all bees share broadly similar diets (mainly nectar and pollen), the social species—particularly the corbiculate Apinae, but also certain carpenter bees [25]—are distinctive in harboring specialized gut bacterial symbionts in the adult stage [17,26]. These symbionts, including *Snodgrassella*, *Gilliamella*, and host-restricted *Lactobacillaceae* and *Bifidobacteriaceae* lineages, appear to have been present throughout much of the evolutionary history of the corbiculate Apinae, though with losses in some bee lineages [27]. Their long-term persistence and specialization may be facilitated by the co-occurrence of adult parents and offspring within a shared nest, a defining feature of eusociality [28], which provides a transmission route for microbes.

Findings from the most diverse, yet least understood group of eusocial bees, stingless bees (Meliponini), show that sociality per se is not sufficient to maintain specialized symbionts. A large survey of the genus *Melipona* [29], and smaller-scale surveys of other taxa [26,30–35] have shown that *Snodgrassella* and *Gilliamella* are absent in some, but not all, stingless bee lineages. The reasons why these bacterial symbionts vary among stingless bee species are unknown but may be related to the particularities of host social structure. Most stingless bees form perennial colonies with a single long-lived queen, a continuous worker caste, and food stores, with new colonies founded through swarming [36]. However, stingless bee species vary substantially in the number of workers that make up the mature colony (i.e., the colony size). For example, *Plebeia minima* colonies may have as few as 100 workers [37], whereas *Trigona amazonensis* can surpass 100,000 [36–38]. Colony size, or group size more generally, is a fundamental aspect of social organization [22]. It can influence traits such as communication, task specialization, immunity, and nest ecology [39–41]. Based on theory and prior empirical work on animal group size, the extreme variation in colony size seen among stingless bees can be expected to influence the microbiome. For example, group size may alter the efficacy of symbiont transmission within and between groups [18], as well as selective pressures (e.g., pathogens) that may influence symbiont maintenance [42,43]. This relationship has not yet been tested at a phylogenetic scale.

In comparison with many host traits used for comparative analyses, the microbiome can fluctuate extensively over time, spatially, and between conspecific individuals [5,44,45]. Such intraspecific variability can obscure macro-evolutionary patterns, and the colony structure of social bees may further amplify intraspecific variability in the microbiome. The magnitude of this variability may itself vary among host taxa and contexts. Still, studies on some Australian stingless bees (*Tetragonula* and *Austroplebeia*) have demonstrated that they maintain host-specific bacterial communities [46] even across large geographic distances [31]. Overall, there remains a need for deep within-species sampling in other clades, particularly the Neotropical species, which make up the large majority of stingless bee diversity.

Here, we reconstruct the evolutionary history of gut symbiont gain and loss in stingless bees, and test whether these dynamics are driven by colony size. To do so, we first sampled a large number of worker bees from five Neotropical stingless bee species (∼20 to ∼80 individuals per species) and characterized their microbiomes using 16S rRNA gene sequencing. This helped us to determine how well a small number of individuals and colonies would capture a bee species’ characteristic microbiome. Then, we used a taxonomically broad collection of stingless bees (workers of 29 species and 18 genera) [47], 16S rRNA gene sequencing, quantitative PCR, and phylogenetic comparative methods to reconstruct gut microbiome dynamics over the course of host diversification. Our findings highlight group size as a key driver of microbiome evolution in stingless bees and, potentially, in social animals more broadly.

## Methods

### Sample collection

From a global stingless bee collection [47], we selected 187 adult worker bees from 51 stingless bee species for microbiome analysis (Table S1). These species were chosen based on the availability of colony size estimates in the literature (Table S2). We used the maximum reported colony size for each species for comparative analysis (described below). Note that methods used to estimate colony size vary between studies and in some cases are approximations. Most of the species we included in our study are Neotropical, with only a few from Africa, Asia, or Australia. No species were collected across multiple continents. Specimens were stored in 95% ethanol, which has been shown to be a suitable buffer for microbiome analysis [48]. Specimens were collected across multiple years and research projects. While some were collected directly from a nest entrance, others were collected from flowers and may represent workers from multiple colonies, and this information was not always recorded due to the opportunistic nature of the sampling. Therefore, colony identity cannot be confirmed for these samples.

Specimens from five additional species were collected at the entrances of their nests in early 2018 from three locations in Costa Rica. Two of these locations are in tropical dry forest at 300 m elevation in Santa Rosa National Park in the province of Guanacaste, separated from one another by approx. 24 km, and one is in montane cloud forest at ∼1400 m elevation in the Monteverde region in Puntarenas (Table S3, Fig. S1). In total, 189 bees were sampled, from the species *Partamona orizabaensis*, *Tetragonisca angustula*, *Tetragona ziegleri*, *Trigona fulviventris*, and *Scaptotrigona subobscuripennis* (Table S3). All specimens were preserved in 95% ethanol and stored at –20 °C.

Each sample analyzed in this study corresponds to an abdomen from a single adult worker bee. Whole abdomens were used to characterize the gut microbiome because, while they contain other tissues, prior work on bees has shown that abdomen or even whole-body homogenates yield microbiome profiles similar to those from dissected guts (e.g., [48,49]). Surface sterilization was not performed. This procedure has been shown not to affect microbiome composition in insects (including honeybees), likely because internal microbial biomass greatly exceeds surface-associated microbial biomass, especially after storage in ethanol [48]. Abdomens were removed from specimens and transferred into bead-beating tubes containing 750 µL ZymoBIOMICS lysis solution and two 5 mm 440C stainless steel grinding balls. Homogenization was carried out in a bead-beater at 1800 rpm for 2 minutes, followed by a 1-minute rest, another 2-minute cycle, and a final 1-minute rest [50]. Homogenates were stored at −70 °C until DNA extraction.

### DNA extraction, library preparation, and sequencing

Lysed samples were used to extract genomic DNA (gDNA) using the ZymoBIOMICS 96 DNA Kit, following the manufacturer’s protocol. Negative extraction controls and Zymo mock community standards were included. Extracted gDNA was PCR-amplified using barcoded universal primers, 515F (5’-GTGCCAGCMGCCGCGGTAA-3’) and 806R (5’-GGACTACHVGGGTWTCTAAT-3’), targeting the V4 region of the 16S rRNA gene [51]. PCR reactions (25 µL) included 12.5 µL Platinum Hot Start PCR Master Mix, 1 µL each of forward and reverse primers (10 µM), and 1 µL of template DNA. The thermal cycling profile consisted of an initial denaturation at 94 °C for 3 min, followed by 35 cycles of denaturation (94 °C, 45 s), annealing (50 °C, 60 s), and extension (72 °C, 90 s), with a final extension at 72 °C for 10 min. PCR products were cleaned and normalized using SequalPrep normalization plates and pooled for sequencing. Amplicons were sequenced on an Illumina MiSeq platform (2 × 250 bp paired-end reads) at the UC Irvine Genomics Research and Technology Hub. Raw sequence data are available in the ENA (accession number PRJEB97714).

### Quantification of total bacterial load

To estimate absolute bacterial abundance per bee abdomen, we performed absolute quantification of 16S rRNA gene copy number using quantitative PCR. Amplification targeted the V1–V2 region of the 16S rRNA gene using universal primers 27F (5’-AGAGTTTGATCCTGGCTCAG-3’) and 355R (5’-CTGCTGCCTCCCGTAGGAGT-3’). Reactions with PowerUp SYBR Green Master Mix (ThermoFisher) were run in 10 µL volumes in triplicate, with 1 µL of template DNA per reaction [50,51].

Standard curves were generated from a 16S rRNA gene standard, serially diluted from 10^8 to 10^2 copies/µL, and included on every plate in triplicate. Each qPCR plate included a no-template control. Amplifications were performed under the following conditions: 95°C for 10 min; five cycles of 95°C for 15 seconds, 65°C for 15 seconds, and 68°C for 20 seconds; then 35 cycles of 95°C for 15 seconds, 60 for 15 seconds, and 68°C for 20 seconds. Absolute copy numbers per bee abdomen were calculated by multiplying the estimated concentration in the genomic DNA (copies/µL) by the elution volume (20 µL), and then by 750/400 (1.875), to account for the fraction of the 750 µL abdomen homogenate carried into the extraction (400 µL, per the ZymoBIOMICS 96 DNA Kit protocol). To estimate the absolute abundance of individual bacterial taxa per bee abdomen, relative abundances (i.e. proportions of 16S rRNA gene amplicon sequences) of each focal taxon were multiplied by the total bacterial load (16S rRNA gene copies per abdomen) for each sample. These per-taxon absolute abundances were then averaged across conspecific individuals for visualization and analyses.

### Amplicon data analysis

Paired-end reads were quality filtered using Trimmomatic v0.38 [52] with a 4-base sliding window and a minimum Phred score of 20. A headcrop of 15 bp and a minimum read length of 180 bp were applied. Denoising, chimera removal, and inference of amplicon sequence variants (ASVs) were performed with DADA2 [53] in QIIME2 v2024.10 [54], truncating forward and reverse reads to 230 bp. Taxonomy was assigned using a Naïve Bayes classifier trained on the SILVA v138 reference database [55] with the q2-feature-classifier plugin in QIIME2 [56]. A total of 2,267,484 reads and 5,847 ASVs were initially recovered, with an average of 11,934 reads per sample before filtering.

Following best practices for sequencing-based insect microbiome studies [57], and using sequence data from our extraction negative controls, we identified and removed putative reagent-derived contaminants using the decontam R package v1.18.0 [58] with the prevalence method (threshold = 0.5). Samples with more than 25% contaminant sequences were flagged as being contaminated and excluded from further analysis. This step reduced the dataset to 29 species and 108 individuals.

Data filtering and visualization were conducted using the phyloseq v2.4-4 [59], MicroViz v0.11.0 [60], ggplot2 v3.5.2 [61] and smplot2 v0.2.5 [62] packages in R. Taxa unassigned at the phylum level or classified as eukaryotes were removed. To remove very rare and potentially spurious ASVs, we applied a minimum per-sample abundance threshold (≥10 reads in at least one sample). After rarefying to 2,500 reads per sample, the final filtered dataset included 617 ASVs and 267,500 reads across 107 samples.

SILVA-derived taxonomic classifications were refined for ASVs belonging to Neisseriaceae and Orbaceae, the families that include the focal genera *Snodgrassella* and *Gilliamella* (respectively). All relatively abundant ASVs (mean relative abundance across samples > 0.1%) classified as Neisseriaceae or Orbaceae were queried against the NCBI nt database using MMseqs2 v15 [63]. For each query, we retained the top 10 highest-scoring hits. Accession numbers from these hits were mapped to NCBI taxonomic identifiers using taxonomizr v0.11.1, and the genus identity of the Lowest Common Ancestor (LCA) for each lineage was inferred using the condenseTaxa function. For one Neisseriaceae ASV in the Costa Rican dataset the LCA result was ambiguous, and so a manual blastn search was conducted to confirm its identity.

Microbial community composition was analyzed using centered log-ratio (CLR) transformation implemented in MicroViz. Principal component analysis (PCA) was performed on genus-level CLR-transformed data, with relative abundances aggregated to genus level by summing read counts across congeneric ASVs, to visualize patterns in community structure. Beta diversity was assessed using Bray–Curtis dissimilarities. To quantify the contribution of each taxon to overall microbiome variation, we calculated the loadings of each genus along the principal components of the PCA. Alpha diversity was estimated for each sample using the Shannon index, calculated with the phyloseq package v2.4-4 [59].

### Phylogenetic analyses

To investigate the evolutionary dynamics of specialized gut symbionts, we used a five-gene Bayesian stingless bee phylogeny from a previous study [47]. Note that the individuals originally sequenced to construct the phylogeny are colony siblings to those sampled for microbiome analysis here. The original tree was pruned to the 29 species included in our microbiome dataset and then transformed into an ultrametric tree using the chronos() function in the ape R package v5.7-1 [64] under a correlated model. We tested several values of the smoothing parameter lambda (0, 0.2, 0.4, 0.6, 0.8). Analyses were consistent across trees, so we retained the tree generated with lambda = 1 for downstream analysis.

Microbial abundances were aggregated to the genus level as described above to obtain relative abundances for the symbionts *Snodgrassella*, *Gilliamella, Lactobacillus*, and *Commensalibacter*. For each bee species, we calculated the mean relative abundance of each symbiont and dichotomized them as present (>0.1%) or absent (≤0.1%). The 0.1% threshold was chosen prior to analysis with the goal of minimizing false positives that may arise from cross-contamination during DNA extraction or index hopping during sequencing.

To infer *Snodgrassella* and *Gilliamella* transitions across the stingless bee phylogeny, we used ancestral state reconstruction with stochastic character mapping via the make.simmap() function in the phytools R package v2.4-4 [59]. Model testing indicated that the equal-rates (ER) model provided the best fit compared to alternatives, so we used it for subsequent analyses. For each symbiont, we estimated 1000 trait histories under an ER model to generate a distribution of ancestral states and transition events across the phylogeny. Uncertainty in ancestral states was quantified as the frequency of each state across the 1000 stochastic character maps, summarized with describe.simmap(), and visualized as pie charts at each node with densityMap().

To test whether the evolutionary histories of *Snodgrassella* and *Gilliamella* gain and loss were correlated, we applied Pagel’s binary trait correlation test (fitPagel()) as implemented in phytools [59]. Here, we assessed whether the trait histories of the two symbionts were independent or dependent of each other using a likelihood ratio test (LRT).

We assessed associations between symbiont prevalence and colony size using two approaches differing in the treatment of traits as discrete versus continuous. First, we implemented phylogenetic generalized least squares (PGLS) regression to model relationships between a species’ maximum reported colony size and either the relative abundance (mean proportion of sequences across conspecific individuals) or absolute abundance (mean estimated number of 16S rRNA gene copies) for each symbiont taxon. These analyses account for the possibility that bee species vary not only in the presence/absence of a symbiont, but also in its abundance (either proportional or absolute) when present. Both relative and absolute abundances were log-transformed before fitting models, as these values spanned multiple orders of magnitude. PGLS models were fitted using the gls() function in the nlme package v3.1-163 [65], with a Brownian motion correlation structure specified via corBrownian() from the ape package. Second, treating both symbiont presence and colony size as binary variables, we conducted binary trait correlation tests as described above. Colony sizes were dichotomized as large or small based on whether they were above or below the median.

Two additional phylogenetic tests were performed. We used a phylogenetic ANOVA to assess whether the absolute abundance of bacteria, as measured by qPCR, differed between species with and without *Snodgrassella*, *Gilliamella* or *Commensalibacter*. To test whether alpha diversity was associated with host colony size, we applied PGLS regression as described above, with log-transformed colony size as the predictor and mean Shannon index as the response variable.

## Results

To assess how intraspecific variation might affect inference of interspecific gut microbiome patterns in our broad phylogenetic survey, we analyzed five stingless bee species in Costa Rica, each represented by 20–80 workers belonging to 1-4 colonies, depending on the species (Fig. S1; Table S3). Microbial community composition in abdomen samples strongly differed among host species (Fig. S2, A; PERMANOVA: *R*² = 0.65, *p* < 0.001). Microbiome composition also differed between colonies in the two species for which we have colony-level replication, *Tetragonisca angustula* (Fig. S2 B; PERMANOVA: R² = 0.18, p < 0.001) and *Partamona orizabaensis* (Fig. S2 C; PERMANOVA: R² = 0.21, p < 0.001), although these effects are much weaker than the species effect. Rarefaction curves of total microbial diversity plateaued after sampling ∼5 workers per species, even when multiple colonies and locations (within Costa Rica) were pooled (Fig. S3). Patterns for the symbionts *Snodgrassella* and *Gilliamella*, the focus of the analyses below, are more variable. In some bee species, one or both symbionts are consistently present in conspecific individuals (Fig. S4). In others, some individuals harbor *Snodgrassella* or *Gilliamella* while other conspecifics do not (i.e., ≤ 0.1% relative abundance) (Fig. S4). For example, in *T. angustula*, each colony contains some bees with *Gilliamella* and some without *Gilliamella* (Fig. S2 B).

To examine patterns and drivers of microbial gain and loss at a macroevolutionary scale, we used 16S rRNA gene sequencing of specimens from a global stingless bee collection [47], with 1–5 workers per species (median = 3). Across these samples, an average of 78% of 16S rRNA reads belonged to five bacterial genera: *Lactobacillus*, *Commensalibacter*, *Gilliamella*, *Snodgrassella*, and *Bifidobacterium*, and to unclassified members of Acetobacteraceae and Lactobacillaceae (Fig. 1). The other 22% belong to a long list of rare taxa, none of which exceed 2% mean abundance. The same taxa were also highly abundant in our five deeply sampled species from Costa Rica (Fig S6).

**Figure 1.**
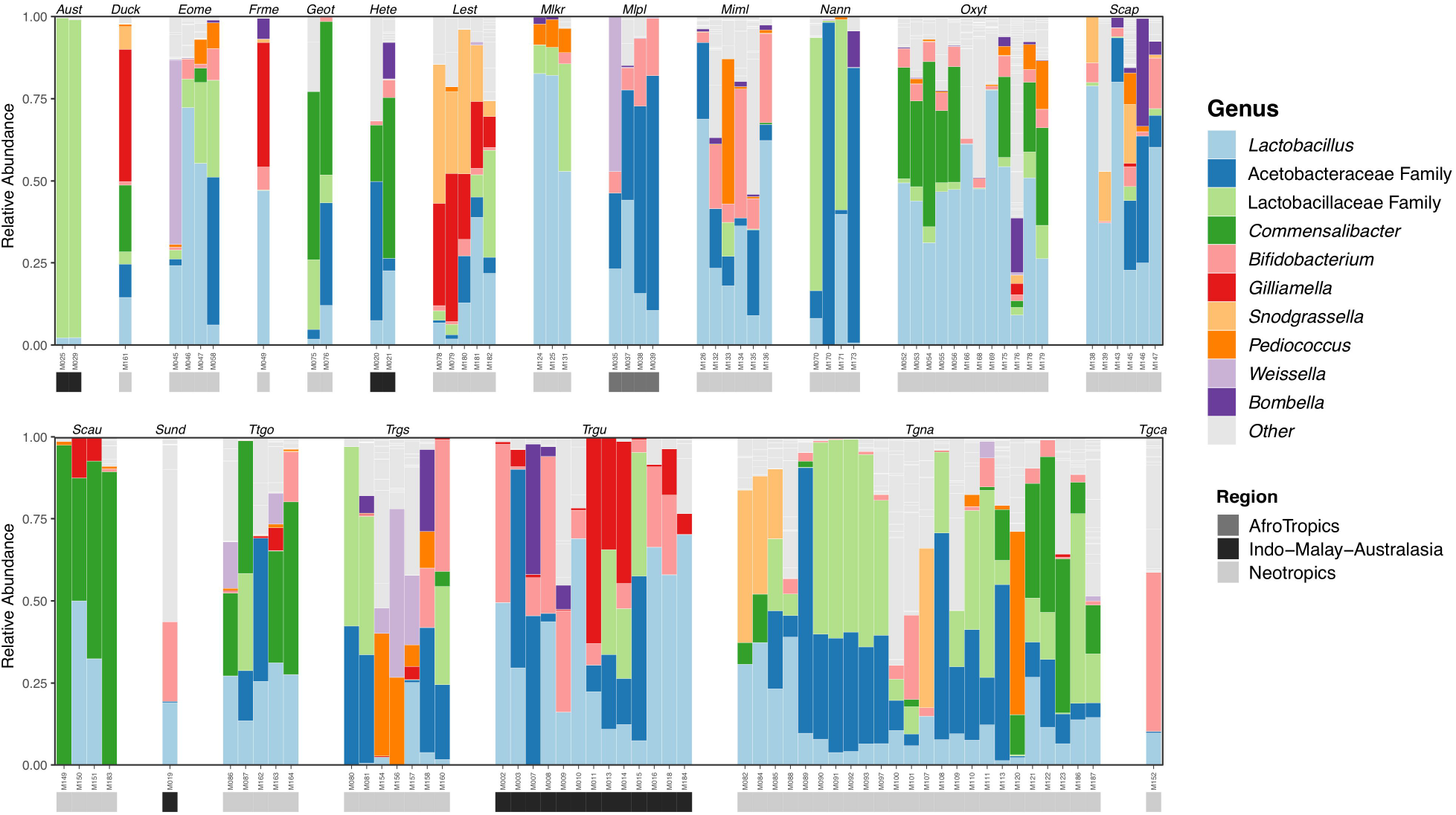
Gut microbiome composition of stingless bees. Each bar corresponds to an individual bee, and the relative abundance of each bacterial genus is represented by different colors. Two bacterial taxa lacking genus-level classifications are listed in the legend with their family-level classifications. The bottom strip represents the regional origin of each bee. Samples were ordered by bee genus. The 10 most abundant bacterial genera are shown (others collapsed as ‘Other’).

The bacterial taxa that contributed most to overall microbiome variation among samples were *Snodgrassella* and *Gilliamella* (Fig. 2), which together accounted for 70.1% of the variance along PC1 (34.9% and 35.2%, respectively), exceeding any other genera. *Lactobacillus* and *Commensalibacter* were also among the most abundant taxa and are included in select analyses below (Fig. 1). Total absolute abundances of bacteria did not systematically differ across species based on the presence or absence of *Snodgrassella* (F = 2.27, p = 0.298; Fig. S8A), *Gilliamella* (F = 0.02, p = 0.929; Fig. S8B) or *Commensalibacter* (F = 0.42, p = 0.643, Fig S8C). This analysis does not apply to *Lactobacillus* as it was present in all species. The four most-common bacterial taxa show wide variation in estimated absolute abundances among bee species, with distinct patterns for each bacterium (Fig. S9).

**Figure 2.**
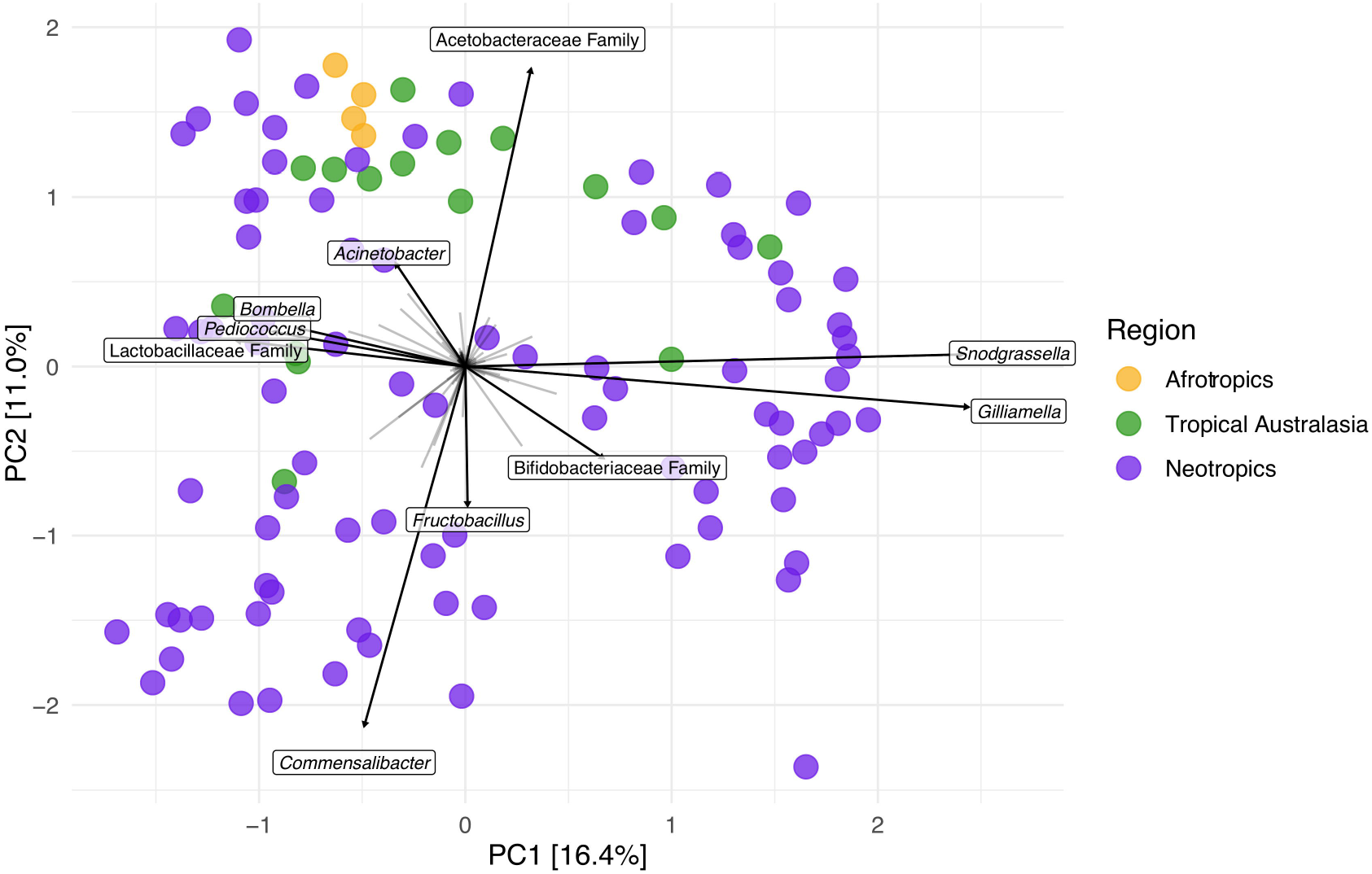
Principal component analysis of center log-transformed bacterial relative abundances at the genus level. Arrows showing the top taxa loadings in the ordination. Taxa that could not be confidently resolved to genus are labeled at the lowest classified rank (family). Each dot represents an individual bee, colored by region.

We used ancestral state reconstruction to infer gains and losses of gut symbionts during stingless bee evolution, focusing on *Snodgrassella* and *Gilliamella*. We recover complex evolutionary histories for both symbionts, each characterized by multiple independent gains and losses (Fig. S7). The most recent common ancestor of Meliponini most likely lacked *Gilliamella*, but whether it did or did not harbor *Snodgrassella* cannot be resolved (Fig. S7). We then fit Pagel’s correlated binary state model to test for coevolutionary dynamics—i.e., whether gains and losses of each symbiont (*Snodgrassella* or *Gilliamella*) were dependent on the presence of the other symbiont (Fig. 4A). The dependent model significantly outperformed the independent model (likelihood-ratio test, LRT = 18.20, p = 0.0011), in support of correlated evolution of the two symbionts. The inferred transition rates were asymmetric: *Snodgrassella* can be gained and lost independently of *Gilliamella*, but the acquisition and maintenance of *Gilliamella* appear to be dependent on *Snodgrassella* (Fig. 4A). The presence of *Gilliamella* without *Snodgrassella* is highly unstable, with exceedingly low transition rates to this state (q = 0), and high rates away from it. Indeed, none of the stingless bee species included in this study harbor *Gilliamella* without *Snodgrassella* (Fig. 3A).

**Figure 3.**
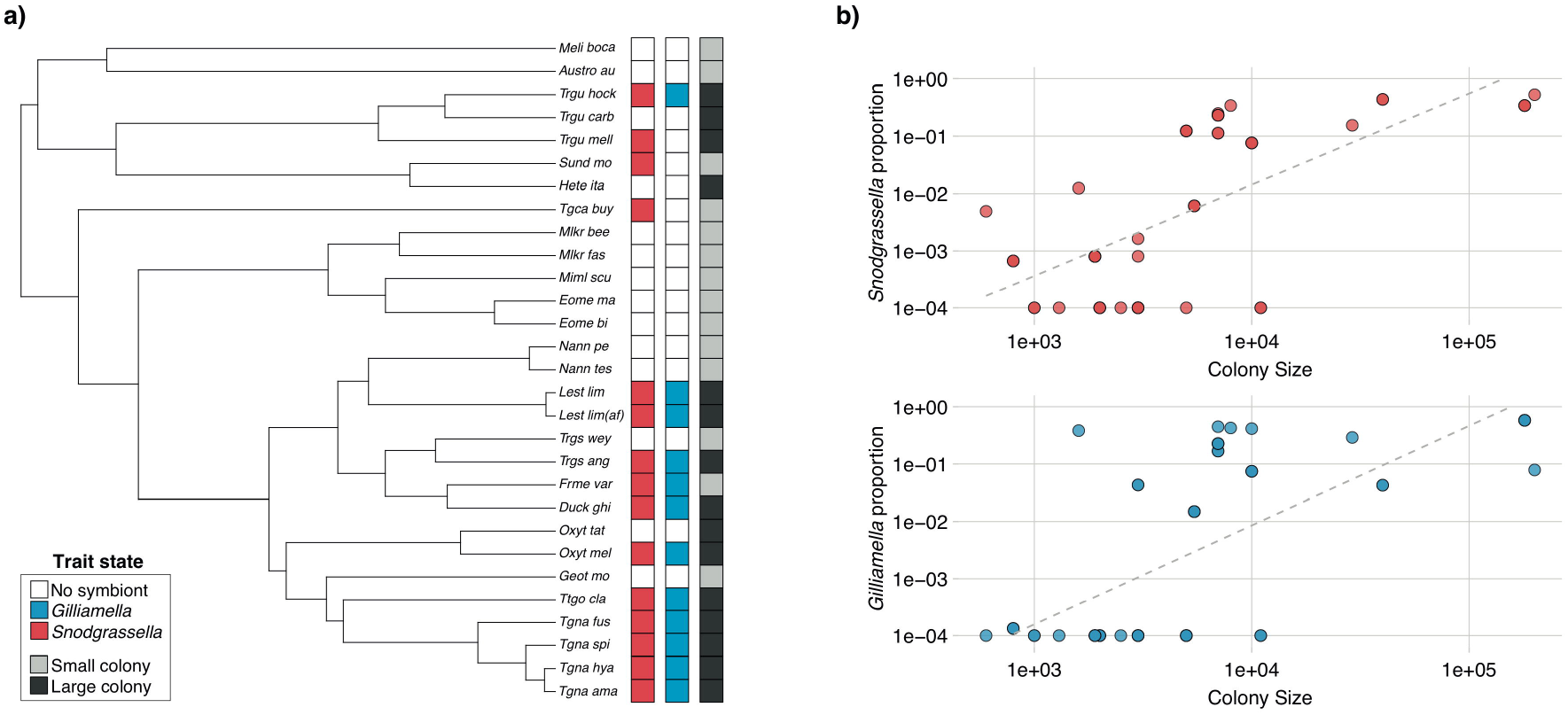
(a) Phylogenetic distribution of *Snodgrassella* and *Gilliamella* symbiont presence across 29 stingless bee species. In the left and middle column, colored boxes indicate the presence or absence of each bacterial group. The right column indicates the species’ maximum colony size, categorized as small (below the median) or large (above the median). (b) Phylogenetic generalized least squares (PGLS) regression between colony size and the species-averaged relative abundance of *Snodgrassella* (top) and *Gilliamella* (bottom). *Snodgrassella* is positively associated with colony size (R² = 0.48, P < 0.037), while *Gilliamella* is not (R² = 0.41, P = 0.077). Dashed lines represent ordinary least-squares regressions plotted for visualization. Points aligned on the x axis correspond to species with 0% prevalence, which could not be log-transformed.

**Figure 4.**
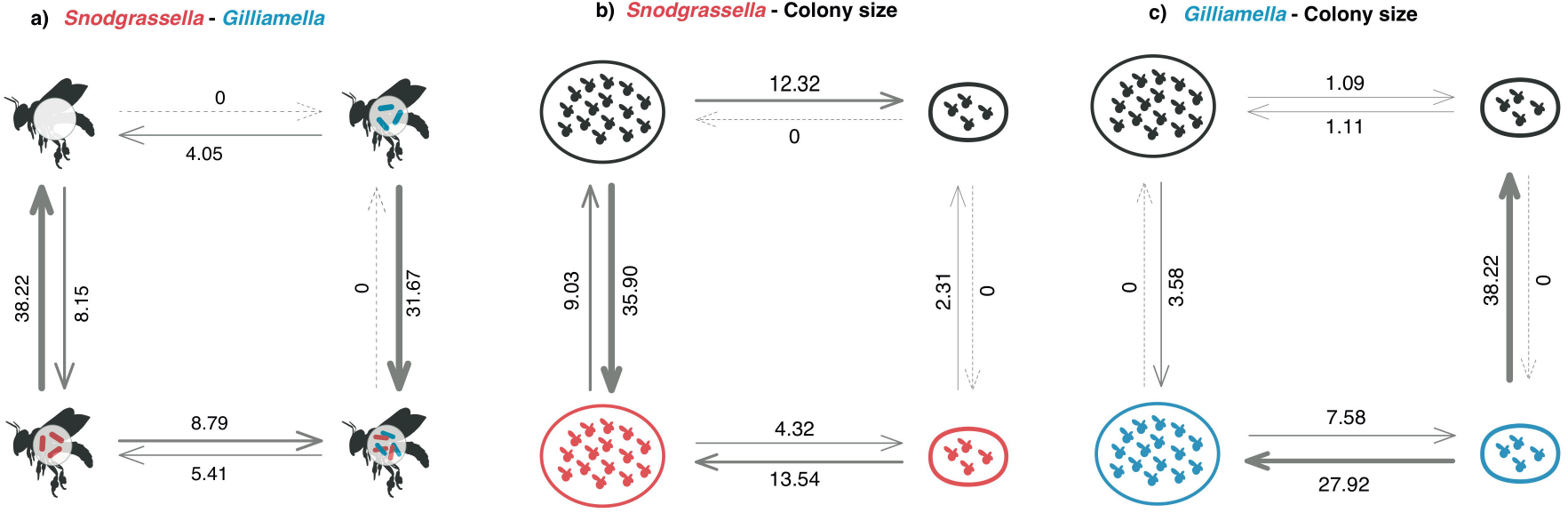
Binary-trait correlations and transition rates from Pagel’s discrete model. The corners of each diagram represent combinations of binary states for the two symbionts (present/absent × present/absent). Thicker arrows correspond to higher transition rates, and arrow labels represent the estimated transition rates. (a) Coevolutionary association between the presence/absence of *Snodgrassella* (red) and *Gilliamella* (blue); the dependent model fits better than the independent model (likelihood-ratio test, LR = 18.2, P = 0.0011). (b) Association between colony size (small vs. large) and *Snodgrassella* presence (LR = 12.8, P = 0.0126). (c) Association between colony size and *Gilliamella* presence (LR = 16.1, P = 0.0028). Arrows show maximum-likelihood transition rates (q□□ ; line width ∝ rate); dashed arrows indicate rates estimated as 0.

Next, we used phylogenetic comparative methods to test associations between host colony size and symbionts. PGLS regressions of symbiont relative abundance on log-colony size showed a strong positive relationship for *Snodgrassella* (R² = 0.48, *p* = 0.037; Fig. 3B) but not for *Gilliamella* (R² = 0.41, *p* = 0.077; Fig. 3B). Conversely, when performing the same analysis but using per-taxon absolute abundances, *Gilliamella* was positively associated with colony size (PGLS: R² = 0.39, p = 0.007; Fig. S10B), but not *Snodgrassella* (R² = 0.49, p = 0.392; Fig. S10A).

To further examine these relationships. we performed a complementary test using Pagel’s correlated-traits model, for which both variables are treated as binary. This analysis showed significant associations between colony size (large versus small) and the presence or absence of each symbiont (likelihood-ratio tests, *Snodgrassella*: LR = 12.75, p = 0.0126, Fig. 4B; *Gilliamella*: LR = 16.14, p = 0.0028, Fig. 4C). For *Snodgrassella*, bee lineages are inferred to evolve large colonies only in the presence of the symbiont (transition rate, q = 13.54), and not without it (q = 0). Similarly, evolutionary transitions to small colonies are less likely in lineages with *Snodgrassella* (q = 4.32) than those without it (q = 12.32). Conversely, the model also indicates that gains and losses of *Snodgrassella* are modulated by colony size. *Snodgrassella* is inferred to be exclusively acquired in lineages with large colonies (q = 35.9). The small-colony-size, no-*Snodgrassella* state appears to be an evolutionary dead end (q = 0 for transitions to alternate states). The *Gilliamella* model also supports a relationship between colony size and symbiont status. As with *Snodgrassella*, evolution of large colonies is much more likely in bee lineages with *Gilliamella* (q = 27.92) than those without it (q = 1.11). Gains of *Gilliamella* only occur in lineages with large colonies (q = 3.58), while losses only occur in those with small colonies (q = 38.22).

Finally, we examined whether other taxa besides *Snodgrassella* and *Gilliamella* covaried with colony size across bee species. Microbiome alpha diversity (mean Shannon index across conspecific bees) was not significantly associated with host colony size (PGLS: β = -0.09, p = 0.213). The relative abundance of *Lactobacillus*, the most common bacterial genus among stingless bees (Fig. 1), was negatively associated with colony size (PGLS: p = 0.002, R² = 0.11). However, this pattern was not significant when using absolute abundance estimates (p = 0.162) (Fig. S10) and, given the compositional nature of the amplicon data, might be due to negative covariance with *Snodgrassella* and *Gilliamella*. *Commensalibacter*, another common bacterial genus (Fig. 1), showed no significant association with colony size using either metric (relative abundance: p = 0.738; absolute abundance: p = 0.774; Fig. S10).

## Discussion

We found that much of the variation in gut microbiome composition among stingless bee species is accounted for by the host-specialized symbionts *Snodgrassella* and *Gilliamella*. These symbionts are widespread (though not universal) across honey bees and bumble bees [26,66,67], as may be expected from symbionts that provide beneficial functions to hosts.

However, our data show that the evolutionary history of these symbionts in stingless bees is highly dynamic. Ancestral state reconstruction indicates that the common ancestor of stingless bees likely lacked *Gilliamella* but could not confidently resolve whether it harbored *Snodgrassella*. As stingless bees diversified, each symbiont was gained and lost multiple times. These findings contradict a previous model in which specialized symbionts were acquired by the common ancestor of corbiculate Apinae bees and then irreversibly lost in some stingless bee lineages [26]. Further, the observation that one or both symbionts are missing from so many stingless bee species raises questions about their fundamental importance to social bee biology. The heterogeneous distribution of the symbionts suggests that their costs and benefits may be context-dependent, varying with the particular biology of their hosts. It should be noted that some bee species may harbor a particular symbiont only in certain parts of their range or during certain times of year. Thus, greater intraspecific sampling might reveal finer-scale microbiome dynamics than we are currently able to observe (see below for more discussion of this issue).

Why some stingless bee species harbor the host-specialized gut symbionts *Snodgrassella* and *Gilliamella*, while others do not, has been a mystery. Here, using phylogenetic models, we found that colony size (the number of bees inhabiting a nest) is linked to the gain and loss of *Snodgrassella* and *Gilliamella* at macroevolutionary scales. In one model we employed, where both variables were treated as continuous, the relative abundance of *Snodgrassella* and the estimated absolute abundance of *Gilliamella*, but not the converse, showed statistically significant increases with colony size. Non-linear models might be useful in clarifying these relationships. In another model where both variables were binary, the relationship between symbiont presence and colony size was significant for both *Snodgrassella* and *Gilliamella*. In the latter model, both symbionts are inferred to have been acquired exclusively in lineages with large colonies; for *Gilliamella*, losses are restricted to lineages with small colonies. Causality remains ambiguous here. For example, *Snodgrassella* cannot be a prerequisite for bees to evolve large colonies, while also being unable to first establish in small colonies, as could be interpreted from the model results. We suggest that symbiont dynamics may both influence, and be influenced by, colony size evolution.

Why might colony size influence the gain and loss of symbionts? One potential mechanism is demographic stochasticity [68]. All else being equal, a microbial population will be more likely to stochastically drift to extinction when it is smaller. Such conditions should occur in bee species with smaller colonies, and hence, smaller collective microbial populations. Another way of viewing this relationship is through the lens of host-parasite models. Decreased host population density generally decreases microbial transmission rates, increasing the probability of microbial extinction [69–73]. Colony size-dependent environmental parameters within the nest may also influence whether symbionts are maintained. For example, small social insect colonies often regulate nest temperature less effectively than large colonies [74–76], and in bumble bees, *Snodgrassella* fails to colonize newly emerged hosts when the temperature is too low [77]. Note that, in theory, the processes described above could operate even if the symbionts are not beneficial to the host.

An alternative explanation for the link between specialized gut symbionts and colony size is that the selective advantage of symbionts is stronger in the context of a large colony. In our transition rate models, the presence of these symbionts is associated with an increased probability of evolving larger colonies. While there is limited experimental evidence of symbiont functions in stingless bees, work on honey bees and bumble bees shows that they may contribute to host pathogen defense and nutrition [78–81]. As explained below, these functions could help hosts overcome the challenges associated with living in large groups.

Parasites and pathogens are generally more prevalent in animals living in larger groups [82], and this disease pressure may favor acquisition and retention of defensive symbionts. In stingless bees, larger colonies conduct more total foraging trips daily [38]. Because they are central-place foragers, where workers leave the nest to collect resources and return to the same location, colonies with more foragers have a greater chance of acquiring a pathogen from the environment. Once acquired, the pathogen is expected to have higher transmission rates due to higher host population density (mentioned above). These processes should also increase the likelihood of superinfection, acquisition of a second pathogen that competes with the first, which may select for greater pathogen virulence [83,84]. Empirical evidence from bees and other insects shows that antimicrobial defenses tend to be stronger in larger groups, indicating that group size can select for enhanced endogenous immunity [85–87]. By the same logic, protective symbionts, which provide a complementary form of antimicrobial defense, could also be favored under these conditions. In honey bees and bumble bees, specialized gut symbionts including *Snodgrassella* and *Gilliamella* have been experimentally shown to reduce pathogen infection [42,78,88]. Symbiont-provided pathogen defense may be especially important for stingless bees; queens are typically monandrous (mating only once), resulting in low genetic diversity within colonies and potentially increased disease susceptibility relative to highly polyandrous species like honey bees [89,90].

Colony size-dependent nutritional demands may also select for maintenance of specialized gut symbionts. In stingless bees, workers forage for pollen and nectar, both for their own consumption and for larvae, and produce glandular secretions and trophic eggs to feed the queen [36,91]. The efficiency of these processes may be disproportionately important for large colonies, for various reasons [92]. For example, workers of large colonies may need to fly farther per foraging trip because they more quickly deplete nearby resources [93,94]. In theory, greater digestive efficiency could support these processes, enabling growth of large colonies. Evidence from honey bees shows that gut symbionts can enhance digestion: *Gilliamella* degrades complex polysaccharides from pollen and monosaccharides from nectar that the host cannot digest [95–97], and the fermentation of these breakdown products results in short-chain fatty acids that may be metabolized by the bee [80].

Gains and losses of the two focal symbionts, *Snodgrassella* and *Gilliamella*, are correlated across the stingless bee phylogeny. However, the dynamics are asymmetric: *Snodgrassella* was capable of persisting in bee lineages independently, whereas *Gilliamella* was never found in the absence of *Snodgrassella*. These patterns suggest that *Snodgrassella* may facilitate *Gilliamella* acquisition and retention, potentially by modulating host immunity [88], metabolite cross-feeding [98] or altering the physicochemical environment (e.g., oxygen) [80]. This facilitative relationship should be tested in experiments with stingless bees.

The patterns of interspecific microbiome variation discussed above are partially supported, but also complicated, by our analysis of intraspecific microbiome variation. In five stingless bee species that were sampled more intensively, colony effects on microbiome composition were weak relative to species effects, and rarefaction curves of microbiome diversity plateaued after sampling ∼5 individual worker bees. Thus, sampling a large number of replicate colonies and individuals may not be necessary to detect interspecific differences in the microbiome. However, it should be noted that the focal species were all sampled within northwestern Costa Rica, and broader sampling could recover alternative patterns. Indeed, previous studies have found variation in stingless bee gut microbiomes between habitats and over time [32,99], which could influence interspecific patterns. Moreover, some of the Costa Rican species showed intraspecific variability in the presence of *Snodgrassella* and/or *Gilliamella*. Our analysis uses a 0.1% mean (across conspecific samples) relative abundance cutoff to score a symbiont as present for a given bee species, which provides some robustness to intraspecific variability. But there is still the potential for false negatives in cases where sample sizes are low. We recommend that future studies perform, for a focal set of stingless bees varying in colony size, deep sampling across the range of each species.

Other caveats should also be taken into account. Some samples had to be excluded from the dataset due to contamination; these samples contained, on average, two–three orders of magnitude lower bacterial biomass (Fig. S5), a factor which increases contamination risk [57]. Further, only a fraction of the ∼550 species of stingless bees [36] are included here, with a particular lack of species from Africa and Asia/Australasia. This, in addition to the lack of range-wide sampling within each species, limits our ability to assess geographic variation in the microbiome. Stingless bee species also vary in numerous ways (beyond colony size) that could matter for the microbiome, such as in nesting ecology, collection of resins, or dietary preferences [36]. If these traits covary with colony size, they may be the direct drivers of symbiont gain and loss. However, such information is not currently available for most species. We hope that this work helps motivate further research on the ecology and life history of stingless bees, which remain poorly understood relative to honeybees and bumblebees.

Whether colony size and host-specialized symbionts covary in other eusocial bees has not been tested with phylogenetic comparative methods, but current evidence suggests a different relationship than in stingless bees. Among honeybees, *Snodgrassella* and/or *Gilliamella* are less prevalent in *Apis florea* and *A. dorsata* than *A. cerana* and *A. mellifera* [26,67] *A. florea* and *A. cerana* have smaller colonies (several thousand bees) than *A. dorsata* and *A. mellifera* (tens of thousands) [100]. Bumblebees have small colonies relative to most stingless bees, yet workers of most, if not all species appear to harbor *Snodgrassella* and *Gilliamella* for at least part of the colony life cycle (seasonal losses are common) [26,101]. Clearly, there are additional factors beyond colony size that influence symbiont maintenance in these taxa.

## Conclusion

Our findings reveal that group size, a fundamental property of all social organisms, is tightly linked with the gain and loss of host-specialized bee gut symbionts over evolutionary time scales. We propose that larger social groups face greater disease risk and perhaps higher nutritional demands, causing greater selective pressure on hosts to recruit symbionts that help them meet those challenges. Larger groups should also provide conditions conducive to symbiont maintenance and specialization, as they harbor larger microbial populations that are less likely to stochastically fail to be transmitted from one host generation to the next. Thus, a feedback loop may arise in which group size and symbiosis reinforce one another. Given the ubiquity of sociality across animals (and other organisms), such feedbacks may operate across a wide range of taxa.

## Data accessibility

Raw 16S rRNA gene amplicon sequence reads for both datasets are available from the European Nucleotide Archive under study accession PRJEB97714. Curated metadata, colony size estimates with their literature sources, ASVs tables with taxonomic assignments, quantitative PCR measurements of total bacterial load, and the host phylogeny are available from the Dryad Digital Repository: https://doi.org/10.5061/dryad.0k6djhbhk. Analysis code for both the global and the Costa Rica datasets is available at https://github.com/nickole97/SB_Pipeline.

## Acknowledgements

We thank Santa Rosa National Park and El Sistema Nacional de Áreas de Conservación (SINAC) for permission to collect stingless bees in Costa Rica (#R-SINAC-ACG-PI-033-2018). Sydney Cameron and Tommy McElrath for loaning specimens, Kristal Watrous for laboratory support, and Luis Alberto Chica and Alberto Barrón-Sandoval for advice on bioinformatic analysis.

